# Sulbactam-mediated inhibition of TEM-1 β-lactamase increases specific yields of ampicillin-resistant plasmids

**DOI:** 10.1101/2025.10.19.683307

**Authors:** Stanislav Pepeliaev

## Abstract

Ampicillin resistance is the most widely used selection marker in molecular biology. Most ampicillin-resistant plasmids originate from highly active TEM-1 β-lactamase cassette which causes very fast antibiotic decay and negatively affects long-term plasmid retention. The present work proposes a method of controlling the β-lactamase activity by the addition of an irreversible β-lactamase inhibitor. The positive effect of a β-lactamase inhibitor, specifically sulbactam, on plasmid retention is confirmed by much higher specific plasmid yields: 4-8 times higher titres in shake-flask and almost 3 times higher in bioreactor scales were observed. Among all commercially available β-lactamase inhibitors sulbactam provides the best combination of affordability and activity. The sulbactam addition is simple, fast and economic solution allowing to screen thousands of legacy pBR322 or pUC-derived plasmids harbouring TEM-1 β-lactamase cassette without engineering their sequence.

## Introduction

Last decade is characterised by an increased interest to seemingly well-known biology and chemistry of plasmids. This has connection to the increased demand in plasmid quantities for gene therapies (Tiscornia, Singer and Verma, 2006; Segura *et al*., 2013; Emmerling *et al*., 2016; Merten, 2016; Merten, Hebben and Bovolenta, 2016), transient and cell-free protein expression(Steger *et al*., 2015; Melinek *et al*., 2023) and vaccine production (Pascolo, 2007; Williams, Carnes and Hodgson, 2009; Scott *et al*., 2015; Liu, 2019). In other words, plasmids have become a staple in biotechnology, which in turn puts a lot of pressure on economics of plasmid production.

Main traits of plasmids enabling their easy manufacture is the ability to self-replicate in bacteria and a very simple and efficient selection of bacteria based on antibiotic-resistance cassettes. Historically, the most wide-spread and frequently used selection marker in plasmids is based on ampicillin resistance. More specifically, it is based on the TEM-1 β-lactamase first discovered in the sixties and then wide-spread after becoming the part of pBR322 plasmid. The recent analysis (Cumming *et al*., 2022) has demonstrated that almost all commercially available plasmids with ampicillin resistance use the same native TEM-1 cassette or minor modifications of it. The overall ratio of Amp-resistant plasmids is also very high – the estimated number of Google Scholar citations is above half-a-million (Cumming *et al*., 2022); and among more than 800 plasmids from the Jenner Institute lab collection above 85% are based on TEM-1 β-lactamase cassette. This cassette is reasonably compact at approx. 1100 bp and it provides a convenient way of fast selection of transformed cells by expressing high levels of β-lactamase and giving a growth advantage to the resistant cells. Being among the first discovered and characterised antibiotic resistance cassette, the TEM-1 historically became a default choice of a selection marker for majority of newly designed plasmids.

The mechanism of β-lactamase action is based on the hydrolysis of β-lactam ring, which irreversibly destroys the molecules of an antibiotic (Majiduddin, Materon and Palzkill, 2002). High activity of TEM-1 β-lactamase cassette causes extremely short half-life time of β-lactam antibiotics in liquid media with resistant bacterial cultures: approximately 30 minutes for 100 mg/L ampicillin when inoculated with a single colony and even shorter 6 minutes for a 1:100 diluted overnight culture (Cumming *et al*., 2022). The numbers for the same concentration of carbenicillin were marginally better with the half-life of 45-50 minutes when inoculated with a single colony (Cumming *et al*., 2022). Such a rapid decay of antibiotics severely undermines mid-and long-term plasmid retention. This is especially true for high-copy plasmids: in absence of selective pressure the subpopulations with lower plasmid copy number grow faster rapidly becoming dominant in the bacterial culture. This is not an issue for typical small-scale isolation protocols yielding micrograms of plasmids but becomes a serious obstacle if quantities above tens of milligrams are required, or a long-term plasmid retention is required (Balbas and Bolivar, 1990; Makrides, 1996).

There were several approaches addressing the low plasmid retention caused by excessively high activity of TEM-1 β-lactamase: antibiotic resistance cassette replacement (Jeffrey and Joachim, 1991; Manna *et al*., 2013), downregulation of TEM-1 cassette promoter (Lartigue *et al*., 2002; Takahashi and Sakamoto, 2018; Cumming *et al*., 2022), switching to antibiotic-free selection(Cranenburgh *et al*., 2001; Guo *et al*., 2024; Marie and Scherman, 2024). All approaches require replacement of certain features of the target plasmid by means of molecular biology. The option of replacing the ampicillin resistance with another, for example kanamycin resistance is among the most frequent, especially taking into the account the use of ampicillin in vaccine production is not welcomed by regulatory agencies due to likelihood of hyper allergic reaction to it in patients (Smith, 1994). Both antibiotic-resistance cassette replacement and switching to antibiotic-free selection require significant changes to the original plasmid design. The mutagenesis of the promoter sequence of TEM-1 cassette (Cumming *et al*., 2022) is less invasive and can be easily achieved with commercially available mutagenesis kits. However, the mutagenesis of tens and hundreds of legacy plasmids would require a significant amount of time and efforts. Most of commercially available mutagenesis kits (Agilent QuickChange Lightning Site-Directed Mutagenesis Kit; Thermo Scientific Phusion™ Site-Directed Mutagenesis Kit; NEB Q5® Site-Directed Mutagenesis Kit) are PCR based. These kits can introduce 1-50 bp changes (insertions, deletions, replacements) into plasmid sequences by synthesising the whole new plasmid from the existing original template. The usage of high-fidelity polymerases requires specifically designed primers and reaction conditions. The newly generated plasmid DNA is then transferred to competent cells and the selection process on agar plates is following. Subsequently, the new forms of plasmid must be propagated and isolated from the single colonies, followed by either Sanger or whole-plasmid sequencing to verify the success of mutagenesis. Larger plasmid rearrangements, such as excision or insertion of larger fragments would require even more work and time even if advanced techniques such as NEB NEBuilder® HiFi DNA Assembly of Gibson assembly reactions are used. Considering the mutagenesis process runs without complications it will still require at least one week of laboratory work and will cost several hundred dollars in reagents, consumables and analytics.

Whenever there is a need in a quick and cost-efficient screening of hundreds of legacy plasmids in plasmid-hungry applications the mutagenesis costs might become a significant obstacle. A simpler solution would allow to quickly screen and sort plasmid focussing mutagenesis efforts only on a small number of high-performing candidate molecules. The presented study proposes such a solution. It is based on a simple method of controlling β-lactamase activity by adding an irreversible β-lactamase inhibitor (clavulanic acid, tazobactam, sulbactam, etc) together with ampicillin or carbenicillin. The simultaneous addition of a lactam antibiotic and a β-lactamase inhibitor, specifically sulbactam, to the bacterial culture increases the half-life of the antibiotic and assures better selective pressure providing better plasmid yields in shake flasks and stirred-tank bioreactors.

## Materials and Methods

### Reagents and consumables

The fermentations in shake flasks and bioreactor were performed using either Gibco LB medium 1x or Bacto CD Supreme FPM medium (Bacto CD), both were purchased from Thermo Fisher Scientific. Additional reagents for medium preparation and bioreactor feed medium: glycerol, magnesium chloride hexahydrate, ammonium hydroxide, thiamine hydrochloride – were purchased from Merck. Carbenicillin, clavulanic acid and sulbactam used for β-lactamase suppression experiments were purchased from Merck or Fisher Scientific.

### Bacterial strain and plasmids

Escherichia coli strain NEB 5-alpha C2987H (New England Biolabs) was used throughout this study. Two high-copy-number plasmids ADP395 and ADP709 were provided by Douglas Group of the Jenner Institute, University of Oxford. Plasmid ADP395 is 3945-bp NanoLuc reporter vector pNL1.3.CMV from Promega (GenBank Accession Number JQ513378). Plasmid ADP709 is 8220-bp derivative of pTwist-CMV-oriP vector carrying the gene of anti-SARS-CoV-2 immunoglobulin heavy chain variable region and woodchuck hepatitis virus posttranscriptional regulatory element.

### Plasmid concentration assay

To determine specific plasmid yield (Y_S_) the culture sample corresponding to 2 mL of OD_600_=1 was taken and processed using QIAprep Spin Miniprep Kit (Qiagen) according to the manufacturer’s manual. To minimise the plasmid losses and to increase the measurement accuracy the plasmid was eluted into 200 µl. The purified plasmid concentration was measured using NanoDrop One (Thermo Fisher Scientific) and the obtained value in ng/µl (equivalent to mg/L) was converted to mg·L^-1^·OD^-1^.

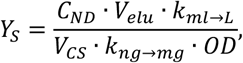

*C*_*ND*_ – plasmid concentration measured by NanoDrop, ng/µl;

*V*_*elu*_ – elution volume, 200 µl;

*k*_*ml→L*_ – conversion from mL toL, 1000;

*V*_*CS*_ – volume of culture sample, mL;

*k*_*ng→mg*_ – conversion of ng to mg, 1000000;

*OD* – bacterial culture density at 600 nm, 1.

Using the following values the formula of specific plasmid yield simplifies to:

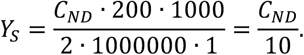

The volumetric plasmid yield (Y_P_) is defined as specific plasmid yield multiplied by OD_600_ and is measured in mg·L^-1^:

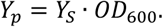

### Plasmid copy number estimation

Depending on the medium and physiological state of bacterial culture the cell size and therefore the cell number varies at the same optical density. Ratio between optical density and cell number varies between 0.5·10^9^ and 1.5·10^9^ cell/ml at OD_600_=1 (Sezonov, Joseleau-Petit and D’Ari, 2007; Wang *et al*., 2009; Sutton, 2011; Couto *et al*., 2018; Beal *et al*., 2020). For this study the ratio OD_600_ of 1.0 = 0.8·10^9^ cells/ml was selected as the most frequently mentioned value. The plasmid copy number (PCN) then was calculated as follows:

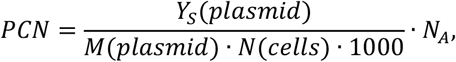

*PCN*– plasmid copy number, copies/cell;

*Y*_*S*_ *(plasmid)* – specific plasmid yield, mg·L^-1^·OD^-1^;

*M(plasmid)* – plasmid molar weight, g/mol;

*N(cells)* – number of E. coli cells in 1L of a culture with OD_600_=1, for this study it is 8·10^11^ cell/L;

*1000* – conversion from mg to g;

*N*_*A*_ – Avogadro number, 6.02·10^23^ molecules/mol.

The molar weights of the plasmids were calculated from their sequences: M(ADP395) = 2437591 g/mol; M(ADP709) = 5079135 g/mol. By substituting constants with their numerical values, the formula is simplified to

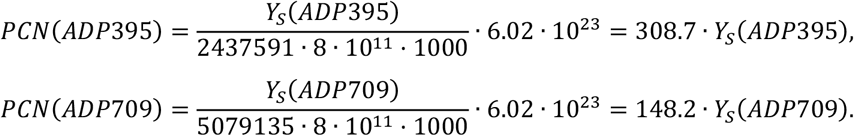

### Shake flask fermentation

Shake-flask experiments were performed in 250-500 mL single-use flasks with baffled bottom and vent caps (Repligene) in 50-100 mL of either LB or Bacto CD medium in a shaking incubator set to 37°C and 225 rpm. The cultivations were performed in presence of 100 or 200 mg/L carbenicillin. The concentration and time of addition of both clavulanic acid and sulbactam varied in wide ranges depending on the experimental design.

The initial shake-flask experiments with clavulanic acid were performed in LB medium at 37°C using plasmid ADP709. The flask with 100 mL LB medium was inoculated with the overnight starter culture to 2% v/v. After reaching the OD_600_ >2 the culture was diluted with fresh medium to OD_600_ = 1 and split into several 50-mL tubes with 10 mL of culture in each; clavulanic acid was added to each sample to the final concentration between 0-100 mg/L. The cultures were incubated in presence of clavulanic acid for one hour, then 100 mg/L of carbenicillin was added to each tube. The fermentation continued for additional six hours. Over the next six hours the cell density and growth rate of each sample was recorded.

The plasmid retention dependence on clavulanic acid concentration was tested using both LB and Bacto CD media. The fresh LB medium was inoculated with 2% overnight inoculum and then grown to OD_600_>2. The OD was adjusted to 2 and then split into three tubes. Straight after 20 and 60 mg/L of clavulanic acid were added to two tubes, the third tube was used as a control. 100 mg/L of carbenicillin addition followed one hour later. For the next three hours the OD_600_ was monitored, and at the end the biomass was collected by centrifugation and used for plasmid extraction using Qiagen mini kit.

The Bacto CD starter culture was grown overnight at 37°C in presence of 100 mg/L carbenicillin. The culture was spun down; the cells were quickly resuspended in fresh Bacto CD medium and split into three 250-mL flasks with 50 ml of culture in each. After 1 hour recovery cultivation clavulanic acid was added to two flasks to 10 and 30 mg·L^-1^·OD^-1^ concentrations; and the third flask was left as a control. One hour later 100 mg/L carbenicillin was added to all three flasks, and the cultivation continued for another three hours. The biomass was collected by centrifugation and used for plasmid extraction using Qiagen mini kit.

The shake-flask experiments with sulbactam were performed in Bacto CD medium with either ADP709 or ADP395 plasmids. The overall layout of experiments replicated those with clavulanic acid with the following differences: the delay between sulbactam and carbenicillin addition was reduced to zero; sulbactam concentrations always were relative to OD_600_ of the culture and varied between 0 and 820 mg·L^-1^·OD^-1^; carbenicillin concentration was either 100 or 200 mg/L.

### Bioreactor fermentation

The bioreactor experiments were performed in Sartorius Biostat B system equipped with 10-L autoclavable glass vessel with starting volume of 3L of Bacto CD medium supplemented with 15 g/L glycerol. Dissolved oxygen (DO) was maintained at 30% by airflow and agitation cascade throughout the fermentation, the temperature was kept at constant 37°C and the pH was maintained at 7 by addition of 25-28% ammonium hydroxide. Carbenicillin and sulbactam were added to the bioreactor prior to inoculation to 200 mg/L and 50 mg/L respectively.

The bioreactor was inoculated with 100 mL of the overnight culture grown in Bacto CD at 37°C in presence of 200 mg/L carbenicillin and 50 mg/L sulbactam (corresponding to 150-250 mg·L^-1^·OD^-1^). The starter culture density threshold was set to OD_600_>8 so that the relative sulbactam concentration in the bioreactor after inoculation is withing 150-250 mg·L^-1^·OD^-1^.

After the depletion of glycerol in the medium detected by the rapid increase of dissolved oxygen (DO spike), the exponential feeding was initiated with 60% w/v glycerol, 16 g/L MgCl_2_·6H_2_O, 0.4 g/L thiamine hydrochloride. The feed rate was programmed to maintain the growth rate µ=0.14 over approximately 7-9 hours. The growth rate µ is defined as follows:

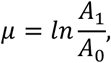

*A*_*0*_ – initial bacterial culture density measured as absorbance at 600 nm,

*A*_*1*_ – bacterial culture density after one hour of fermentation.

The exponential feeding set to maintain specific growth rate means that every hour of fermentation the feed rate increases *e*^*µ*^ times, or 1.15027 times for µ=0.14. The feed rate was recalculated every program cycle or every 5 seconds assuring stepless feed rate increase.

## Results

### Clavulanic acid shake flask experiments

The results of clavulanic acid inhibiting concentration testing are shown on Figure 1. The biomass accumulation (Figure 1A) and growth rate (Figure 1B) trends reveal the cells respond the most to the clavulanic acid presence between first and second hour after carbenicillin addition (second and third hour of the experiment). Apart from the sample with 100 mg/L clavulanic acid all other samples were able to recover from the stress and demonstrated stable growth in later hours of fermentation. In absence of any data points between 30 and 100 mg/L it is safe to conclude the inhibitory concentration of clavulanic acid lays between these two values.

**Figure 1.**
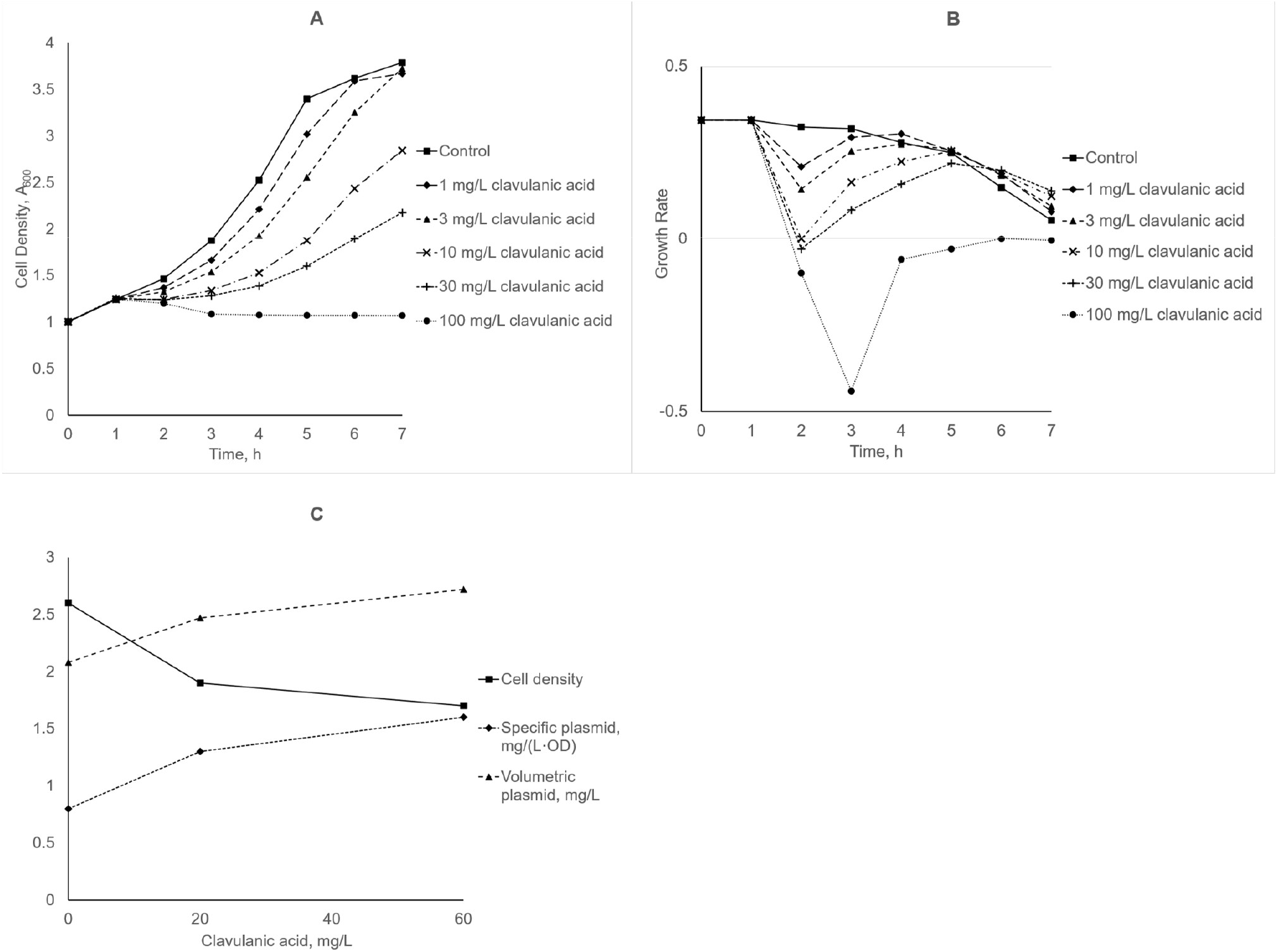
β-lactamase inhibition experiments with clavulanic acid. **A.** Suppresion of cell growth in LB medium with increasing concentrations of clavulanic acid. The highest tested clavulanic acid concentration 100 mg/L led to complete cell growth inhibition. **B**. Time course of growth rates depending on the concentration of clavulanic acid. Note sharp decrease in growth rate between first and second hours after the addition of clavulanic acid. **C**. Impact of clavulanic acid concentration on biomass production (OD_600_), specific and volumetric plasmid yield after six hours of cultivation.

For testing the impact of clavulanic acid on plasmid retention two concentrations were chosen: 20 and 60 mg/L. The Figure1C shows the final cell densities, specific and volumetric plasmid yields. The addition of carbenicillin slowed down the growth rate in all three samples, but more so in presence of clavulanic acid. The sample with the highest clavulanic acid concentration had signs of partial lysis: increased viscosity of the medium and particles of cell debris aggregates were observed. However, after two hours all cultures had recovered and demonstrated further growth. The plasmid retention represented by specific plasmid yield Y_S_ was proportional to clavulanic acid concentration: 0.8; 1.35 and 1.6 mg·L^-1^·OD^-1^ for 0, 20 and 60 mg/L clavulanic acid, respectively. This corresponds to PCNs of 120; 200 and 240 copies per cell.

Bacto CD medium feasibility for high-cell-density cultivation pursuing higher volumetric plasmid yields, was assessed in subsequent experiment. The same absolute clavulanic acid concentrations 20 and 60 mg/L didn’t show any significant difference from control, likely due to higher cell density. Second attempt took into account higher biomass accumulation in Bacto CD medium. The clavulanic acid concentrations were scaled to the cell density. The overnight starter culture reached OD_600_=16. After resuspension in fresh medium and splitting into three flasks the initial cell density was OD_600_=8. The cells were left to recover for 1 hour, then clavulanic acid was added to two flasks to 10 and 30 mg·L^-1^·OD^-1^ – corresponding to relative concentrations in LB medium experiment, with OD_600_=2 at the time of addition of clavulanic acid. The absolute concentrations were 100 and 300 mg/L reflecting cell density OD_600_=10. The specific plasmid yield trends remained similar to LB experiment: 0.96; 1.5 and 1.55 mg·L^-1^·OD^-1^ for 0, 10 and 30 mg·L^-1^·OD^-1^ clavulanic acid, respectively. The final biomass in Bacto CD was much greater reaching OD_600_ between 15 and 18 and delivering 4-5 times higher volumetric plasmid yields.

### Sulbactam shake flask experiments

The shake flasks experiments were continued with sulbactam as a β-lactamase inhibitor and Bacto CD Supreme FPM (Thermo Fisher Scientific) medium as a growth medium. The plasmids used in these experiments were ADP709 and ADP395. Both plasmids behaved identically in experiments, and the choice of a plasmid was based on laboratory needs. Two key changes were made during these tests based on previous and new findings.

First, the simultaneous addition of carbenicillin and β-lactamase inhibitor increases inhibitory effect compared to the delayed addition of carbenicillin. Moreover, this greatly simplified the experimental design, therefore, simultaneous addition of sulbactam and carbenicillin was used for all subsequent experiments.

Second, the activity of β-lactamase inhibitor is biomass-dependent and more dense cultures require proportionally higher concentrations of the inhibitor to achieve the same levels of plasmid retention. Sulbactam experiments demonstrated trends similar to clavulanic acid series. Thus, in the culture with initial OD_600_=0.65 and 130 mg/L sulbactam the specific plasmid yield reached 4.45 mg·L^-1^·OD^-1^ after six hours of cultivation (PCN = 1370 copies per cell). In contrast the culture with initial OD_600_=9 and 135 mg/L sulbactam reached only 0.81mg·L^-1^·OD^-1^ (PCN = 250 copies per cell). The absolute sulbactam concentrations in both examples are nearly identical, the concentrations relative to the biomass differ significantly: 200 mg·L^-1^·OD^-1^ in low-density culture and 15 mg·L^-1^·OD^-1^ in high-density culture. Therefore, in all following experiments the sulbactam concentrations were relative to the cell density.

The relative inhibiting concentration of sulbactam was determined (Figure 2) by testing the range of sulbactam concentration between 0 and 820 mg·L^-1^·OD^-1^. The experiment was carried out in presence of 100 mg/L carbenicillin at 37°C for six hours. The growth rate falls below zero and stays negative in samples with more than 205 mg·L^-1^·OD^-1^. For the plasmid retention experiments the sulbactam concentrations up to 250 mg·L^-1^·OD^-1^ were selected and tested (Figure 2C). The conditions were the same: 100 mg/L carbenicillin was used, the temperature was set to 37°C and the duration of cultivation was six hours. The highest values of specific plasmid yields accompanied by moderate biomass accumulation were observed between 100 and 200 mg·L^-1^·OD^-1^ of sulbactam. At 250 mg·L^-1^·OD^-1^ a significant drop in both cell density and plasmid specific yield was registered. In absence of sulbactam the specific plasmid yield was 0.6 mg·L^-1^·OD^-1^ (PCN = 190 copies per cell). The addition of 200 mg·L^-1^·OD^-1^ sulbactam led to 7-8-fold increase of specific plasmid yield reaching 4.5 mg·L^-1^·OD^-1^ (PCN = 1400 copies per cell).

**Figure 2.**
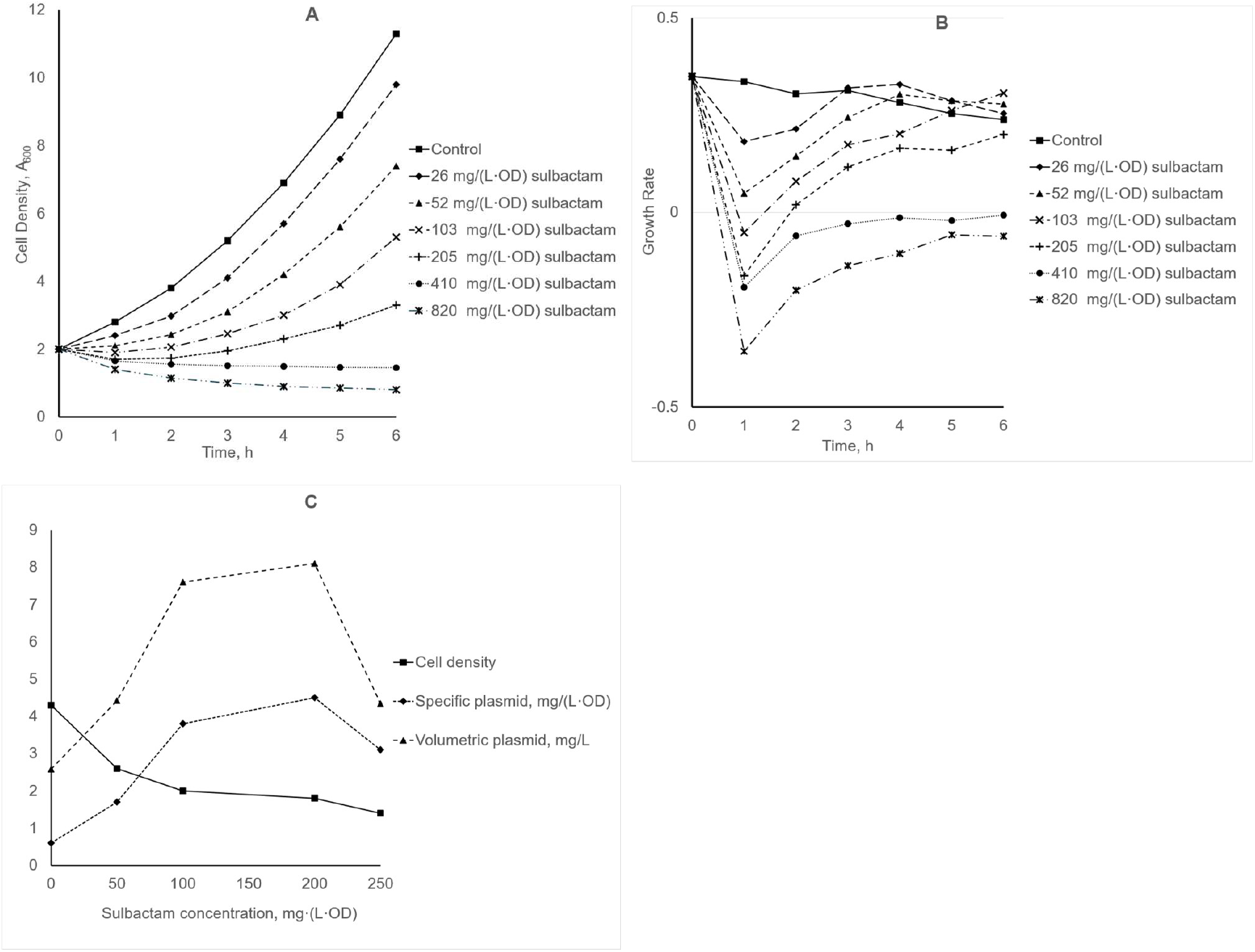
β-lactamase inhibition experiments with sulbactam. A. Suppresion of cell growth in Bacto CD medium with increasing concentrations of sulbactam. The concentrations avobe 400 mg·L^-1^·OD^-1^ fully inhibit cell growth. B. Impact of increasing sulbactam concentration on growth rate. The growth rate drop is proportional to the amount of added sulbactam. C. Impact of sulbactam concentration on biomass production (OD_600_), specific and volumetric plasmid yield after six hours of cultivation.

To fully utilise the potential of Bacto CD medium the longer cultivation times of 20-24 h are recommended by the manufacturer. In presence of 100-200 mg/L carbenicillin and without sulbactam the final biomass commonly reached OD_600_ >24. Unfortunately, longer fermentations are detrimental for the specific yield: it falls from 0.6 to 0.5-0.55 mg·L^-1^·OD^-1^ (PCN = 150 copies per cell). The addition of 50 mg/L sulbactam to the culture with the initial OD_600_ 0.2-0.3 (corresponding to relative concentration 133-200 mg·L^-1^·OD^-1^) was tested in presence of 100 or 200 mg/L carbenicillin (Table 1). This resulted in 4-5 times higher specific and 2 times higher volumetric plasmid yields in Bacto CD and even eight times higher volumetric yields if compared to standard LB cultivations (Table 1).

**Table 1.** Comparison of effects of carbenicilin concentration and sulbactam addition in extended overnight cultivation in LB and Bacto CD media.

|  | Carbenicillin,<br>mg/L | Final<br>OD<sub>600</sub> | Specific<br>plasmid yield,<br>mg·L<sup>-1</sup>·OD<sup>-1</sup> | Plasmid copy<br>number,<br>copies/cell | Volumetric<br>plasmid yield,<br>mg/L |
| --- | --- | --- | --- | --- | --- |
| LB control | 100 | 4.2 | 0.7 | 215 | 2.9 |
| Bacto CD control | 100 | 22 | 0.5 | 150 | 11 |
|  | 200 | 21 | 0.55 | 165 | 11.6 |
| Bacto CD<br>sulbactam 50 mg/L | 100 | 8.8 | 2.3 | 710 | 20.2 |
|  | 200 | 8.7 | 2.7 | 835 | 23.5 |

### Sulbactam stirred-tank bioreactor experiments

To assess the scalability of sulbactam application to bench-top bioreactor scale two fermentations were performed in 10-L stirred-tank bioreactor. The first run without sulbactam addition to both starter culture and bioreactor served as a reference. During the second run 50 mg/L of sulbactam was added to both starter culture (corresponding to approx. 170 mg·L^-1^·OD^-1^ relative concentration) and bioreactor (corresponding to approx. 100 mg·L^-1^·OD^-1^) simultaneously with 200 mg/L carbenicillin. The fermentations started with 15 g/L glycerol in the medium and after the glycerol depletion the exponential feeding was activated at the speed maintaining the growth rate µ=0.14. The biomass accumulation and changes in specific plasmid yield were monitored over 20-h long runs (Figure 3). In both cases negative changes in specific yields were observed: less pronounced decrease from 0.55 to 0.48 mg·L^-1^·OD^-1^ during the control run; and more prominent drop from 2.55 to 1.3 mg·L^-1^·OD^-1^ during the sulbactam run. Nevertheless, it was still 2.9 times higher compared to the control. The difference in final volumetric yields was slightly smaller: in sulbactam run it reached 119 mg/L, compared to 2.6 times lower 46 mg/L in control run – mostly due to slightly higher biomass in control run. The total upstream yield from the whole working volume was approx. 400 mg for the sulbactam run and approx. 140 mg for the control run.

**Figure 3.**
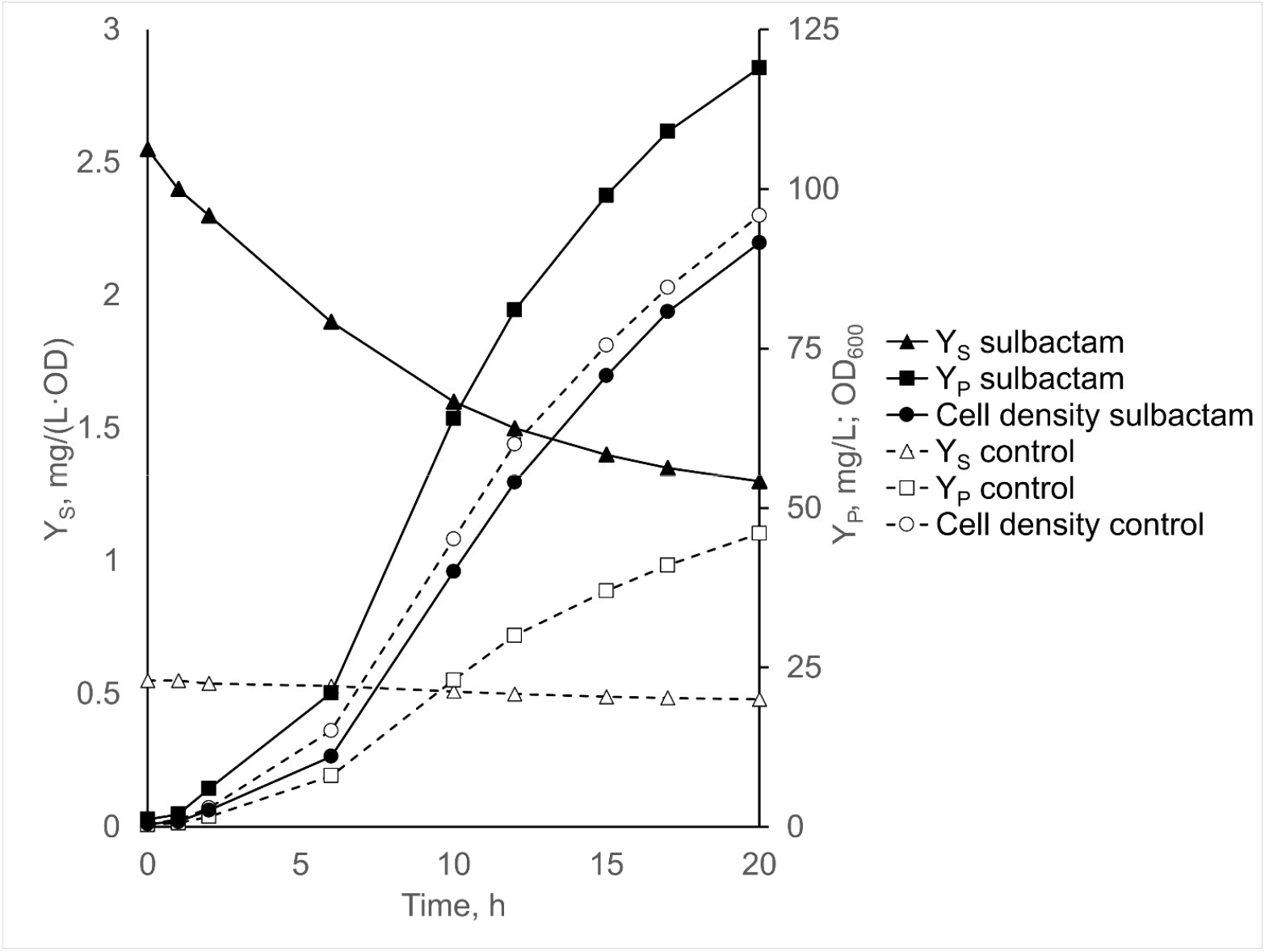
Sulbactam addition effect on specific and volumetric plasmid yields in high-cell-density bioreactor fermentation.

## Discussion

β-Lactamase inhibitors are a group of chemical compounds that are structurally close to β-lactam antibiotics, such as ampicillin or carbenicillin (Figure 4). The chemical similarity, more specifically the presence of a β-lactam ring, allows them to fit into the catalytic centre of β-lactamases and irreversibly inhibit them by binding covalently thanks to the presence of reactive double bonds. The mechanism of reversible inhibition of clavulanic acid differs slightly from that of both sulbactam and tazobactam (Figure 5). However, the irreversible inhibition leads to the same crosslinking of two serine residues in vicinity of catalytic centre.

**Figure 4.**
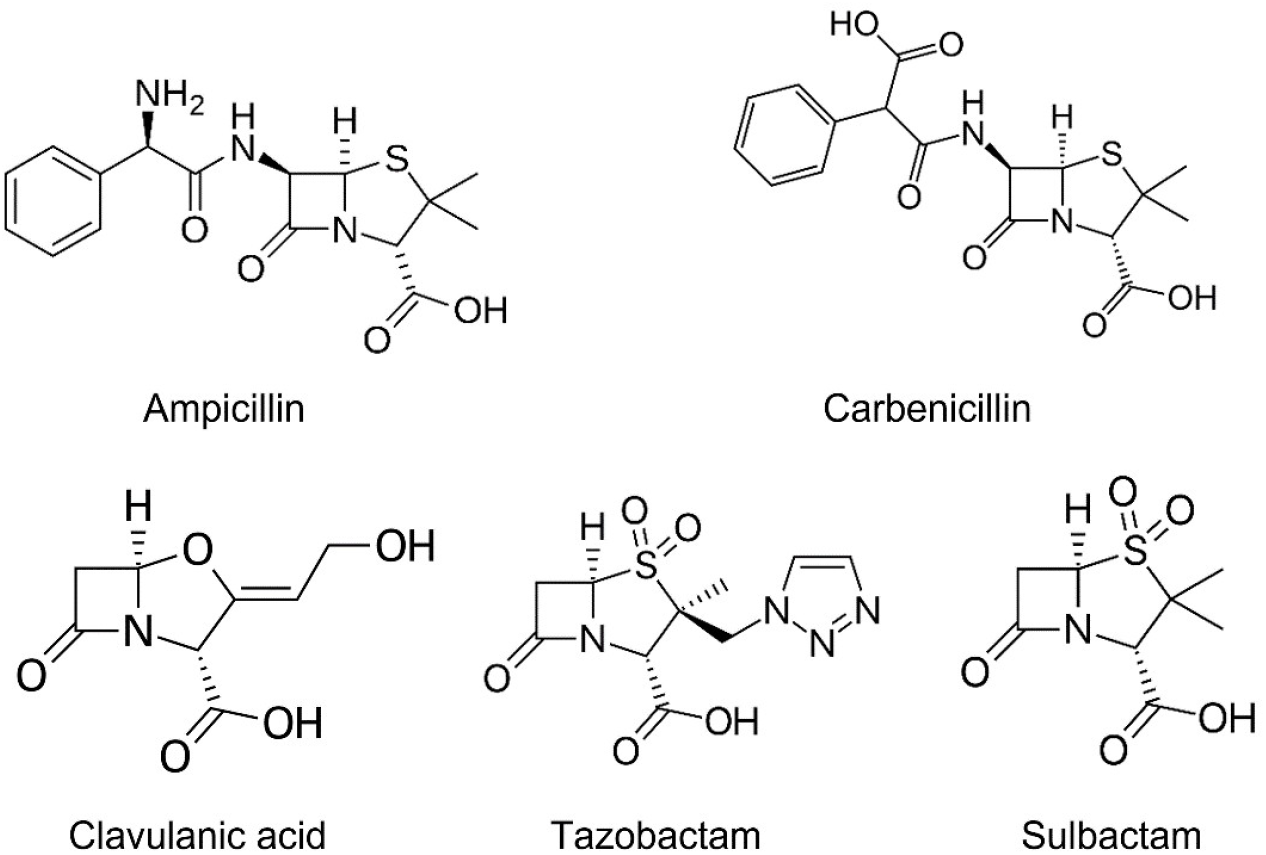
Stuctural formulas of beta lactams: two antibiotics – ampicillin and carbenicillin; and three beta-lactamase inhibitors – clavulanic acid, tazobactam and sulbactam.

**Figure 5.**
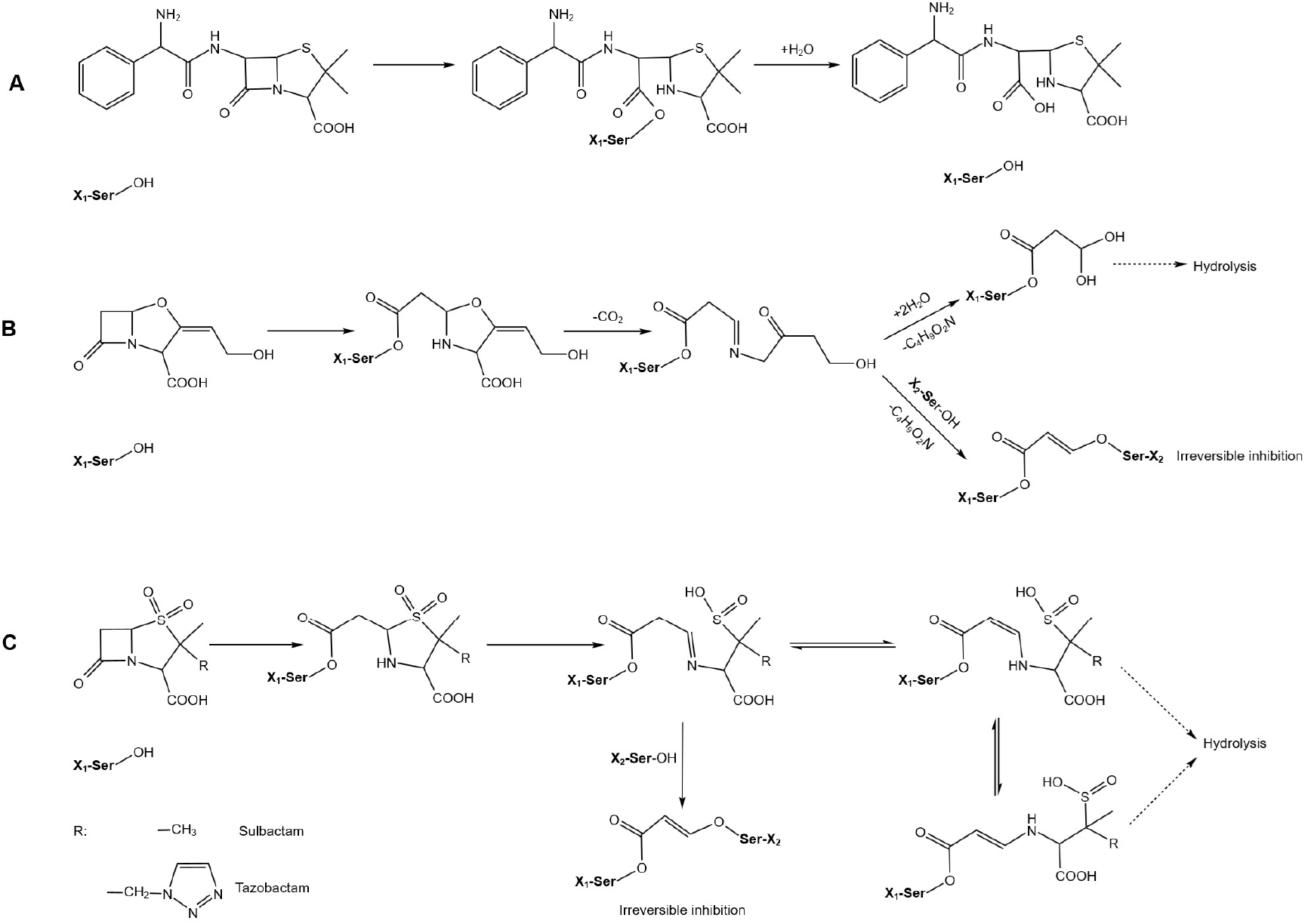
Mechanism of action of β-lactamase and lactams. A. Catalytic hydrolysis of carbenicillin with β-lactamase. B. Scheme of reversible and irreversible β-lactamase inhibition by clavulanic acid. C. Scheme of reversible and irreversible β-lactamase inhibition by sulbactam and tazobactam.

Therefore, the activity of β-lactamase in the medium, cytoplasm and periplasm of the bacterial cells could be controlled by an addition of an appropriate amount of a β-lactamase inhibitor. Such a simple addition of the inhibitor would mimic a weaker promoter and would deliver the same results as promoter region mutagenesis(Cumming *et al*., 2022) in faster and simpler manner and at a fraction of costs of a mutagenesis.

According to the literature analysis the most frequently mentioned and characterised β-lactamase inhibitors are tazobactam (9.2 million Google search results) and clavulanic acid (7.4 million Google search results). The potency of both compounds is similar with tazobactam being slightly superior against certain types of β-lactamases (Payne *et al*., 1994). Clavulanic acid was easier to source and had shorter lead times at the start of the research; therefore, it was selected for the initial studies. First proof-of-concept experiments with clavulanic acid as a β-lactamase inhibitor had shown a principal feasibility of the proposed approach: the addition of a β-lactamase inhibitor led to a decrease of growth rate compared to control. This suggests longer maintenance of higher antibiotic concentration in the culture, which in turn can be possible only if the β-lactamase is inhibited. Simultaneously, as the growth rates decline, the specific plasmid yields increase indicating the average plasmid copy number in population goes up (Figure 1C). It means the presence of β-inhibitor in medium improves selective pressure and discriminates subpopulations with low PCN. This correlates well with the observations of heterogeneous distribution of PCN in bacterial cultures (Jahn *et al*., 2016) indicating the selective pressure is applied by eliminating subpopulations with low plasmid copy numbers.

Given very high price of clavulanic acid and its wide use in clinic it has been decided to switch to much cheaper and more available sulbactam (£1945 per gram of clavulanic acid, T19860-25MG, Merck; £162 per gram of sulbactam, T1631-500MG, Merck). The similarity in mechanism of action of irreversible inhibition (Figure 5) infers β-lactamase inhibitors are mutually replaceable, but differences in activity must be accounted for. In research on β-lactamase inhibitors activities, (Mahoney, Koppel and Turner, 1976; Payne *et al*., 1994; Drawz and Bonomo, 2010; Shapiro, 2017) it was found sulbactam is 10-27 times weaker than clavulanic acid and 11-29 times weaker than tazobactam in inhibiting 35 tested β-lactamases of different origin. More importantly, both clavulanic acid and tazobactam are more efficient in suppressing much broader range of unconventional β-lactamases. As most of the ampicillin-resistant plasmids carry TEM-1 cassette or its derivatives, narrower spectrum and lower activity of sulbactam becomes a safety advantage decreasing the risks of appearance of more resistant mutant forms of β-lactamases. In a very unlikely event of a sulbactam-resistant β-lactamase mutation occurrence such a mutant highly likely will be susceptible to either clavulanic acid or tazobactam. From practical perspective the price difference between clavulanic acid and sulbactam is much greater than the difference in their activities, which makes sulbactam more economical choice.

As the focus of the research was on providing more efficient solution for pDNA production, the LB medium was replaced with chemically defined medium Bacto CD Supreme FPM (Thermo Fisher Scientific). This change brought several benefits. First, higher biomass yields were observed for Bacto CD: the typical biomass density in overnight cultures in LB medium in presence of 200 mg/L carbenicillin was below OD_600_=4, while Bacto CD routinely achieved OD_600_ >20 under the same conditions. Even in presence of high concentrations of sulbactam and carbenicillin the final OD in 22-h overnight culture was reaching OD_600_>8, which is at least two times higher than best results in LB medium. Second, the mineral medium promotes slower growth, which in turn flattens the differences between plasmid-containing and plasmid-free cells improving the plasmid retention(Silva *et al*., 2009). The difference in Y_S_ between control and test samples in LB and Bacto CD confirms this observation: the clavulanic acid addition led to Y_S_ increase from 0.8 to 1.6 mg·L^-1^·OD^-1^ in LB medium and to Y_S_ increase from 0.96 to 1.55 mg·L^-1^·OD^-1^ in Bacto CD medium. Third, a mineral chemically defined medium has a benefit of a better control over the growing culture thanks to a lower number of active metabolic pathways, which is especially beneficial during bioreactor fermentations (Li, Nimtz and Rinas, 2014; Peebo *et al*., 2015). Fourth reason to switch to the chemically defined medium is its GMP compliance, more specifically its animal-free composition. This assures better manufacturability if further scale up or tech transfer is required. Last, but not least, being designed for E. coli fermentations the Bacto CD medium is less likely to support growth of other microbial hosts, therefore lowering chances of contamination and increasing the overall robustness of the bioprocess.

First experiments with sulbactam showed there is a noticeable difference in its activity depending on the time of carbenicillin addition – the longer the delay the lower the sulbactam inhibitory effect. Given the irreversible character of inhibition and very high levels of constitutive expression from TEM-1 cassette(Cumming *et al*., 2022), by the time of antibiotic addition the ratio β-lactamase to sulbactam is higher, causing lesser inhibitory effect. For example, when the same 200 mg·L^-1^·OD^-1^ od sulbactam were used, one hour delay in carbenicillin addition decreased the maximum specific plasmid yield in an overnight culture from 2.7 to 1.52 mg·L^-1^·OD^-1^. The retrospective analysis of clavulanic acid data confirmed this from growth rate perspective: the samples with 2-hour delay between clavulanic acid and carbenicillin addition demonstrated higher growth rates compared to 1-hour delay (Figure 6), which indicates the effective concentration of clavulanic acid decreased over time. Such a behaviour of inhibitor can be explained by the fact that only a small number of their molecules irreversible bind to the enzyme molecule, while the rest is digested. Based on turnover numbers approximately 160 molecules of clavulanic acid and around 10 thousand molecules of sulbactam are required to irreversibly inhibit one molecule of β-lactamase (Drawz and Bonomo, 2010).

**Figure 6.**
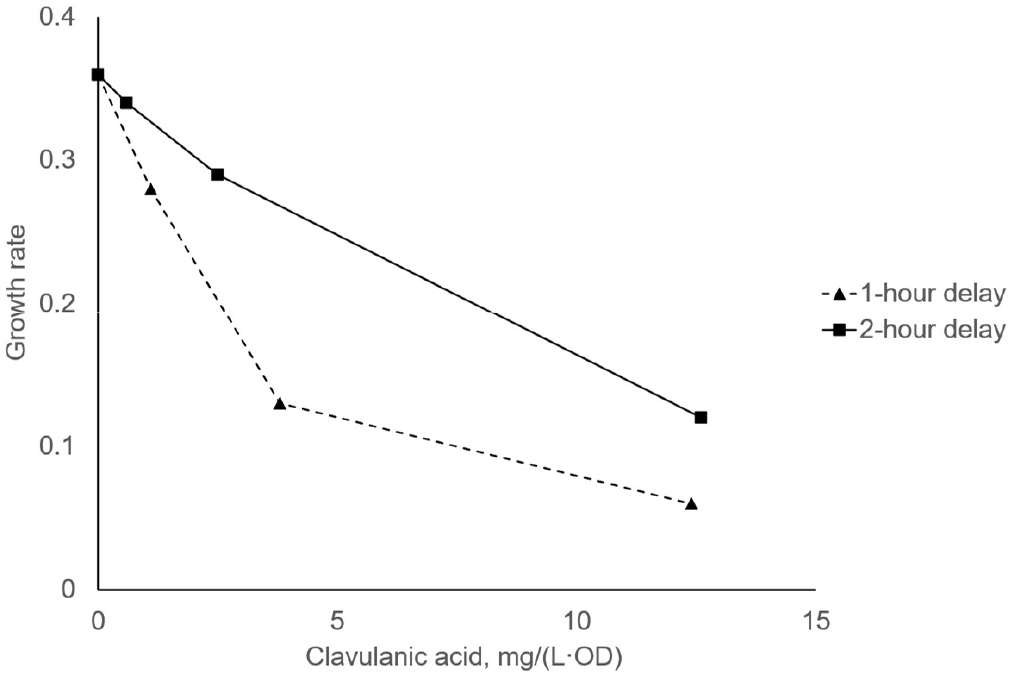
Influence of the delay between carbenicillin and clavulanic acid addition on cell growth inhibition.

The comparison of growth curves shows the concentration of sulbactam required to supress the cell growth is approximately 8-10 times higher than of clavulanic acid. The inhibiting concentration of clavulanic acid is approximately 35-40 mg·L^-1^·OD^-1^ and that of sulbactam is 260-400 mg·L^-1^·OD^-1^. The real difference is likely to be higher, as all the experiments with clavulanic acid were performed with the delayed carbenicillin addition, and this decreases the effect of the inhibitor. Nevertheless, this value is very close to the earlier comparison of activities of different β-lactamases (Mahoney, Koppel and Turner, 1976; Payne *et al*., 1994; Drawz and Bonomo, 2010), where for all tested β-lactamases the activity of clavulanic acid is 10-27 higher than that of sulbactam.

As the present paper focuses on the plasmid production over the protein expression, the measures leading to the highest plasmid yield were prioritised – such as pushing specific and volumetric plasmids yields to the maximum by applying the highest tolerable sublethal amounts of sulbactam and carbenicillin. The application of β-lactamase inhibitor in shake flasks led to an impressive increase of plasmid yields in shake flasks: the volumetric yield increased two-fold when comparing to the control with Bacto CD and eight-fold compared to LB medium (Table 1). The adaptation of antibiotic and inhibitor concentrations may be required if the goal is different: for example, protein expression.

Usually, the shake-flask scale is associated with subsequent plasmid extraction using the commercial kits. All manufacturers of plasmid extraction kits emphasise the criticality of proper ratio between resuspended cells and buffer volumes to assure efficient lysis and neutralisation, and to maximise plasmid yields by utilization of capture columns and membrane filters. Therefore, less biomass with higher specific plasmid yield is preferred over lots of biomass with low specific plasmid yield. A typical Gigaprep kit can deliver 10-25 mg of a plasmid per column by processing 2.5-5 L of an overnight culture grown in LB medium (Qiagen, PureLink, ZymoPure). For example, ZymoPure Gigaprep kit is designed to process up to 20 g of cell pellet to deliver up to 25 mg of plasmid. Therefore, to fully utilise the potential of the kit the specific plasmid yield must be above 1.25 mg/(g WCW). The analysis of the Gigaprep kits from other manufacturers show approximately the same numbers (Table 2). Based on literature (Glazyrina *et al*., 2010; Kangwa *et al*., 2015) one OD unit corresponds to 1.6-1.8 g WCW, in the present study this ratio was found to be 1:1.67. For accurate comparison the WCW values are converted to specific plasmid yields measured in mg·L^-1^·OD^-1^ (Table 2).

**Table 2.** Input material requirements of plasmid isolation kits from different manufacturers.

| Gigaprep kit | Maximum plasmid recovery, mg | Recommended WCW, g | Required specific plasmid yield, $\text{mg}\cdot\text{L}^{-1}\cdot\text{OD}^{-1}$ |
| --- | --- | --- | --- |
| ZymoPure Gigaprep | 25 | <20 | >2.09 |
| Qiagen Plasmid Giga | 10 | <7.5 | >2.22 |
| Macherey-Nagel NucleoBond PC 10000 | 10 | <10 | >1.67 |
| Invitrogen PureLink™ | 12 | <9 | >2.22 |

The minimal specific plasmid yield allowing efficient utilisation of binding columns lay within 1.7-mg·L^-1^·OD^-1^ range for the most popular commercially available plasmid kits. The overnight cultivation in Bacto CD in presence of sulbactam routinely delivers above 2.5 mg·L^-1^·OD^-1^ for the tested plasmid ADP395. Moreover, with further optimisation of growth conditions and concentrations of sulbactam and lactam antibiotic higher values can be expected: the values over 4 mg·L^-1^·OD^-1^ were routinely observed during this study. The results might differ for other plasmids depending on their origin of replication and size, but in general at least two-fold improvement can be expected with the potential of up to eight-fold increase with further process optimisation for a specific plasmid.

Often the scale provided by commercially available kits is not sufficient, or the purity requirements surpass the capabilities of commercially available plasmid isolation kits. In this case scaling up the plasmid process to the bench-top stirred-tank bioreactor or higher is required. Such a step increases the intensity of the bioprocess providing much more input material for further isolation and purification to the specified quality standards. The overall trend favouring less biomass with higher specific yield remains the same, but in this case mostly due to economic considerations. Therefore, a simple method to increase plasmid retention and improve specific and volumetric yields would be welcomed. A simple direct comparison of two bioreactor runs demonstrated the viability of the approach (Figure 3). Even though during the run with sulbactam addition a massive drop of specific plasmid yield was observed from initial 2.55 to 1.3 mg·L^-1^·OD^-1^, the final plasmid yield was 2.9 times higher compared to the control run without the sulbactam addition. Provided a very limited number of bioreactor fermentations and a very simple fermentation strategy there is a high likelihood further process tweaking would assure higher plasmid retention and much higher yields. This could be continuous sulbactam addition with the feed at the rate inhibiting most of β-lactamase activity or lowering the temperature during the batch phase to increase the plasmid stability (Carnes, Hodgson and Williams, 2006). Provided these improvements would be able to maintain the same 2.7 mg·L^-1^·OD^-1^ specific plasmid yield as observed in shake flasks the final volumetric yield would further increase from observed 120 to potential 290 mg/L volumetric yield.

The combination of high-throughput multi-parallel bioreactors (Sartorius Ambr250, Eppendorf DASBox), high-cell density fermentation and optimised sulbactam addition would allow to generate multi-hundred-milligram quantities of several plasmids (up to 24 for Sartorius Ambr250, Eppendorf DASBox) simultaneously without the necessity to replace the antibiotic resistance cassette in each plasmid. This would greatly increase the throughput of testing of many plasmid-hungry therapies and bioprocesses, such as viral vector production, transient expression in mammalian cells or gene therapies. Then, a few best-performing plasmids can undergo the process of targeted mutagenesis significantly reducing associated costs and efforts.

### Sulbactam shake-flask protocol

Based on the findings made during the present work a high-efficiency shake-flask protocol for plasmids with TEM-1 based ampicillin resistance was developed.

1. Transform DH5α or similar strain with the ampicillin-resistant plasmid. Follow the instructions supplied with the competent cells all way down to the incubation on agar plates.
2. Pick a single colony from the agar plate and resuspend it in 2 ml of LB medium with 100 mg/L ampicillin or carbenicillin. Grow overnight in a shaking incubator (200-250 rpm) at 37°C.
3. Prepare 50 ml of Bacto CD Supreme FPM according to manufacturer’s instructions. Add 100 mg/L of ampicillin or carbenicillin and inoculate with 1 ml of the overnight LB culture. Incubate overnight in a shaking incubator (200-250 rpm) at 37°C to adapt the cells to mineral medium.
4. The Bacto CD adaptation is successful if the OD_600_>20 after 20-24 hours of cultivation. If necessary, repeat the cell adaptation to the mineral medium again by repeating the Step 3 and using 1 ml of the overnight Bacto CD culture as inoculum.
5. The fully adapted culture is used for cell bank preparation. The overnight culture in Bacto CD medium with the OD_600_ between 20 and 25 is diluted with sterile glycerol to the final glycerol concentration 20% v/v and OD_600_ = 12-15. The mixture is divided into 2-ml aliquots, and the glycerol stocks are stored at −80°C.
6. Initial sulbactam adaptation is performed by expanding the 2-ml cell bank into 100 ml of Bacto CD medium with 200 mg/L carbenicillin and 50 mg/L sulbactam.
  a. The amount of sulbactam required for a specific plasmid might differ due to different origin of replication, which determines plasmid copy number; plasmid size and toxicity, which contribute to plasmid-replication metabolic burden. For other than pUC-derived plasmids it is recommended to test lower sulbactam concentrations (10-50 mg/L) during the sulbactam adaptation.
  b. The culture is incubated overnight under the same conditions as in Step 3.
7. Depending on the required amount of plasmid the overnight culture from the Step 6 might be processed with the plasmid kit. The expected yields from 100-ml culture with OD_600_=8 and specific yield approx. 2.5 mg·L^-1^·OD^-1^ are 2 mg of the target plasmid. If more plasmid is required, the overnight culture serves as a starter culture for larger volumes by diluting it with fresh Bacto CD medium. The OD_600_ of the diluted culture must be measured to calculate the amount of sulbactam: 100-200 mg·L^-1^·OD^-1^ is recommended for best results. For example, the overnight culture has OD_600_=8. If it is diluted to 1L then the OD_600_ of diluted culture becomes 0.8. Then the amount of added sulbactam should be 80-160 mg/L. For carbenicillin or ampicillin the concentration remains unchanged 200 mg/L. After the dilution the culture is incubated for additional 20-22 hours in a shaking incubator (200-250 rpm) at 37°C.
8. Process culture according to the instructions supplied with the plasmid extraction kit.

Alternatively, the culture from Step 7 can be used as a starter culture for a stirred-tank fermentation. The same rule applies: after the inoculation the OD_600_ of diluted culture must be checked, and sulbactam must be added to the bioreactor to 100-200 mg·L^-1^·OD^-1^ sulbactam.

## Conclusion

The present study addresses the issue of excessively high activity of TEM-1 β-lactamase which leads to a very fact decay of lactam antibiotics in medium and causes low plasmid retention in bacterial cultures. In contrast to other methods the proposed approach doesn’t require plasmid mutagenesis. It relies on the control of β-lactamase activity by addition of an irreversible β-lactamase inhibitor; thus, it is applicable for any plasmid based on TEM-1 cassette. These include numerous plasmids derived from pBR322 and pUC19; plus, many other plasmids sharing the same antibiotic resistance. In the present work 4-kb and 8.2-kb plasmids with pUC origin of replication were used as models and sulbactam was selected as a β-lactamase inhibitor. The effect of β-lactamase inhibitor is time- and biomass-dependent: the longer the delay between β-lactamase inhibitor and carbenicillin addition the lesser the inhibitory effect. Similarly, at the same inhibitor concentration the inhibition is weaker in denser cultures. Both phenomena can be explained by irreversible binding of a finite number of inhibitor molecules to a replenished pool of constitutively expressed molecules of β-lactamase. The higher the number of protein-expressing cells and the longer the reaction time the lesser is the impact of the inhibitor. The best results were observed at the simultaneous addition of sulbactam and carbenicillin to the bacterial culture.

It was demonstrated that the addition of sublethal concentrations of sulbactam leads to a significant increase of specific plasmid yield at cost of lower biomass accumulation. Best results were observed in short experiments, when the bacterial culture was exposed to the combined action of sulbactam and carbenicillin for just six hours: the specific plasmid yield was 7-8 times higher than control meanwhile the biomass was 2.7 times lower (Figure 2C). In more practical scenario of overnight fermentation using chemically defined medium the specific plasmid yield increase was 4.5 times, and the biomass drop was 2.4 times. Finally, in the high cell density fermentation in stirred-tank bioreactor the specific plasmid yield increase was 2.9 times – with just a negligible biomass decline. All these numbers conclude that the suggested approach is viable and potentially applicable for any plasmid carrying a version of TEM-1 β-lactamase cassette.

